# Navigating between global robustness and fine-scale resolution: the potential of demographic inference based on identical-by-descent segments

**DOI:** 10.64898/2026.09.23.752685

**Authors:** Eychenne Océane, Stéphanie Manel, Pierre-Alexandre Gagnaire

## Abstract

Understanding the demographic processes shaping population connectivity is essential for conservation. Yet disentangling the effects of dispersal and local effective density remains challenging because traditional genetic approaches integrate signals over evolutionary timescales. Identity-by-descent (IBD) segments provide opportunities to reconstruct recent demographic connectivity over timescales relevant to contemporary conservation challenges, but their reliability under realistic ecological conditions remains poorly evaluated. Here, we assess the ability of two promising spatially IBD-based methods, *IBD-Analysis* and *MAPS*, to estimate effective density and dispersal variance. Using a simulation–inference framework combining spatially explicit simulations, IBD segment extraction, and demographic inference, we evaluated both methods across a wide range of demographic, ecological and methodological scenarios. Both approaches reliably recovered demographic parameters under homogeneous landscapes, with *IBD-Analysis* providing accurate estimates of global effective density and dispersal across a broad range of conditions. Spatial heterogeneity emerged as the main driver of inference performance. When demographic parameters varied across space, *IBD-Analysis* integrated local variation into biased global estimates, whereas *MAPS* captured spatial patterns but showed substantial variability and generated spurious spatial structure in homogeneous landscapes. Methodological factors had comparatively weaker effects than ecological complexity, highlighting the importance of accounting for landscape structure when applying IBD-based demographic inference. These results highlight the potential of IBD-based approaches to move beyond descriptive patterns of genetic connectivity toward mechanistic inference of demographic processes. By distinguishing the contributions of dispersal and density, these methods can improve conservation decisions by linking genomic patterns to key demographic parameters relevant for conservation.

## 4 | INTRODUCTION

The rapid decline in biodiversity driven by anthropogenic pressures, including climate change, habitat loss and fragmentation, overexploitation, pollution and invasive species, has made understanding the demographic dynamics of natural populations a central challenge for conservation (Barnosky et al., 2011; Hohenlohe et al., 2021; Shaw et al., 2025; Wiens, 2016). Variation in population size and structure underpins long-term population viability by shaping the level and maintenance of genetic diversity (Frankham, 2005; Kardos et al., 2021; Wooldridge et al., 2026), resilience to environmental disturbances (Allendorf et al., 2010; Capdevila et al., 2022) and the risk of local extinction (Lande, 1988; Lowe & Allendorf, 2010; Pérez-Pereira et al., 2022).

Dispersal and local population density are two key determinants of population connectivity. Together, they regulate demographic connectivity (Clobert, 2012) through the balance between immigration and local recruitment (Lowe & Allendorf, 2010) and shape genetic connectivity by modulating gene flow among populations over evolutionary time (Gagnaire, 2020; Gagnaire et al., 2015). Genetic approaches provide complementary insights into connectivity across spatial and temporal scales (Cayuela et al., 2018). Direct methods, such as mark-recapture studies or assignment tests, capture only very recent demographic events, whereas indirect genetic approaches integrate signals over much longer evolutionary timescales (Broquet & Petit, 2009). Consequently, an important methodological gap persists in resolving demographic dynamics over intermediate timescales – spanning tens to hundreds of generations – which correspond to the temporal scale of contemporary anthropogenic changes (Clark et al., 2025; Lande, 1988). At these timescales, disentangling the respective effects of dispersal and genetic drift on spatial genetic structure remains a major challenge.

Spatial genetic structure is classically characterized by increasing genetic differentiation with geographic distance, a pattern known as isolation-by-distance (Wright, 1943). The relationship between genetic relatedness and geographic distance among individuals has been extensively investigated through both theoretical models (Charlesworth, 2009; Hardy & Vekemans, 1999; Robledo-Arnuncio & Rousset, 2010; Rousset, 1997) and empirical studies (Aguillon et al., 2017; Novembre et al., 2008; Puebla et al., 2012; Twyford et al., 2020). In particular, Rousset (1997) showed that the slope of isolation-by-distance is inversely proportional to the product of local effective density (D) and dispersal variance (σ²), highlighting the central role of these two parameters in shaping spatial genetic structure. However, because their effects are confounded, the isolation-by-distance slope alone cannot disentangle the respective contributions of effective density and dispersal, while direct field estimates of these parameters remain challenging to obtain (Cayuela et al., 2018; Puebla et al., 2009). Recovering additional information from molecular data therefore requires moving beyond summary measures of genetic structure. In this context, analyses of genetic relatedness provide a promising alternative by exploiting the recent genealogical relationships among individuals, making it possible to infer the spatial and temporal distribution of shared ancestry.

Recent advances in population genomics reconstruct this genealogical structure by interpreting genetic similarity in terms of shared ancestry explicitly anchored in space. In this framework, genetic relatedness among individuals is viewed as a network of shared ancestors whose topology reflects recent demographic processes (Kelleher et al., 2019a; Kingman, 1982; Wakeley & Sargsyan, 2009). Beyond quantifying the extent of shared ancestry, this perspective reveals where common ancestors were distributed across the landscape and how lineages moved through space over successive generations (Bradburd et al., 2018; Kelleher et al., 2019a), information that has remained largely inaccessible using traditional summary statistics.

Within this genealogical framework, connectivity can be viewed as the spatio-temporal structure of the pedigree linking individuals within a population. Its temporal dimension is determined by the depth of coalescence events: effective population size (N) governs the rate at which lineages coalesce, with smaller populations leading to more recent common ancestors and higher levels of relatedness among individuals (Charlesworth, 2009; Kingman, 1982; Wakeley & Sargsyan, 2009). Its spatial dimension arises from multigenerational dispersal, which shapes the geographic distribution of related individuals by moving lineages across successive generations (Bradburd & Ralph, 2019; Clobert, 2012; Hardy & Vekemans, 1999). Together, these processes determine where and when common ancestors occur within the landscape, giving rise to the observed spatial patterns of shared ancestry and, ultimately, to isolation-by-distance (Bradburd & Ralph, 2019). The recent genealogical structure of a population therefore provides an integrated representation of demographic connectivity across both space and time (Kelleher et al., 2019a; Wohns et al., 2022).

Modern genomic approaches can access this genealogical structure through genomic segments that are identical by descent (IBD), which are inherited from a common ancestor without recombination (Browning & Browning, 2012; Palamara et al., 2012; Sticca et al., 2021). Because recombination progressively breaks down ancestral chromosomes over generations, recently related individuals share longer IBD segments than more distantly related individuals. Consequently, the length of an IBD segment is directly related to the age of the corresponding common ancestor (Browning & Browning, 2012; Palamara et al., 2012), making IBD segments particularly informative about demographic processes occurring over the past tens of generations.

Collectively, IBD segments provide a partial reconstruction of the recent genealogy relating sampled individuals. Combined with the geographic locations of individuals, they provide access not only to the extent of shared ancestry but also to its spatial organization, revealing how lineages have dispersed and coalesced across recent generations (Bradburd & Ralph, 2019; Osmond & Coop, 2021; Ralph & Coop, 2013; Wohns et al., 2022). The joint distribution of IBD segments number, length, and spatial arrangement has thus enabled the reconstruction of recent effective population size (Browning & Browning, 2015; Fournier et al., 2023; Huang et al., 2024), dispersal (Baharian et al., 2016; Ringbauer et al., 2017), and the spatial distribution of recent genetic ancestry (Allentoft et al., 2024; Deraje et al., 2025; Nait Saada et al., 2020; Ralph & Coop, 2013), particularly in human populations.

Building on these properties, a growing number of spatial inference methods have been developed to disentangle the effects of local effective density (D) and dispersal (σ_e_) from the spatial distribution of IBD segments (Al-Asadi et al., 2019; Forien et al., 2024; Ringbauer et al., 2017). Here, we focus on two particularly attractive approaches for molecular ecology studies, *IBD-Analysis* (Ringbauer et al., 2017) and *MAPS* (Al-Asadi et al., 2019), which infer recent demographic connectivity while explicitly incorporating spatial information. However, both methods rely on simplifying assumptions regarding dispersal, reproduction, and underlying demographic models that are rarely met in natural populations (Bradburd & Ralph, 2019; Leblois et al., 2004). Many species exhibit complex life cycles, age structure, non-panmictic reproduction, non-Gaussian dispersal and strong spatial heterogeneity, which may affect inference performance (Clobert, 2012; Hardy & Vekemans, 1999). In addition to these biological assumptions, the methods are subject to data constraints, including accurate haplotype phasing for IBD detection, sufficient sampling, and the availability of informative IBD segments spanning the temporal window of interest (Browning & Browning, 2012). Despite their increasing use (García-Jiménez et al., 2025) and continued methodological development (Campuzano Jiménez et al., 2025; Talbot & Bradburd, 2025), their robustness to realistic biological complexity and methodological limitations has received only limited systematic evaluation.

The objective of this study is to evaluate the conditions under which IBD-based spatial inference can reliably estimate local effective density (D) and effective dispersal variance (σ_e_), therefore providing a robust framework for investigating recent demographic connectivity. To achieve this, we combine forward-in-time spatial simulations incorporating complex life histories and diverse demographic structures, allowing demographic parameters to be controlled while generating realistic genomic data. IBD segments are extracted from simulated recombining genealogies and parameter estimates obtained with *IBD-Analysis* and *MAPS* are compared with the simulated values to assess bias, precision and robustness, thereby delineating the ecological contexts under which these methods provide reliable inference of recent demographic connectivity.

## 5 | METHODS

To assess the accuracy and robustness of demographic inference (dispersal and density) from IBD segments under spatially and biologically explicit scenarios, we implemented a simulation– inference framework (Figure 1). First, we used a spatial genetic model (Figure 1A) to test a range of dispersal and population density values as well as additional demographic and ecological parameters (Figure 1B–C). We then extracted IBD segments from the simulated tree sequences (Figure 1E), while evaluating methodological choices for IBD extraction on demographic inference (Figure 1D). Finally, we compared simulated dispersal and density values with the values estimated by two demographic inference algorithms: *IBD-Analysis* (Ringbauer et al., 2017) and *MAPS* (Al-Asadi et al., 2019).

**Figure 1.**
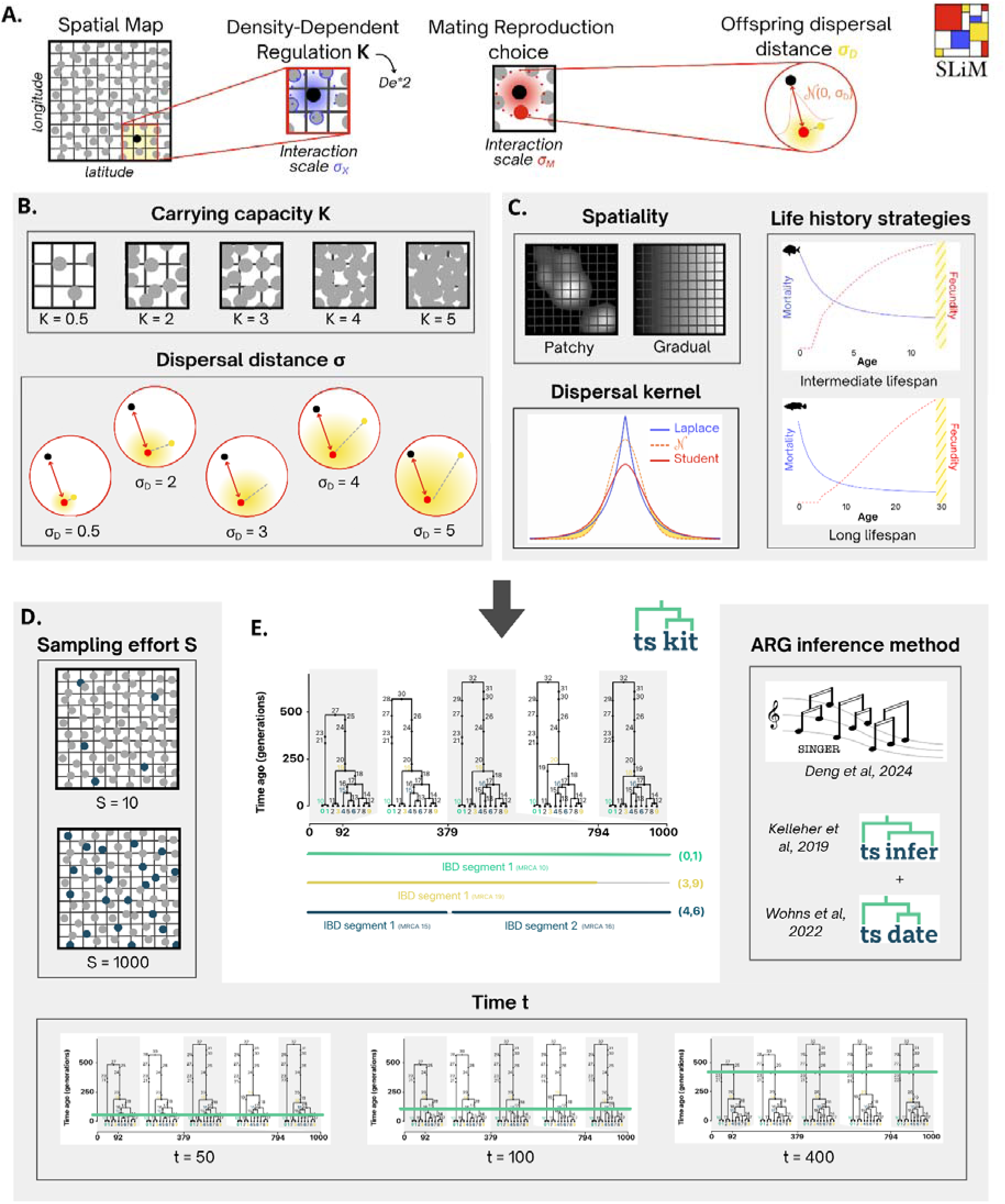
– Simulation and Identity-By-Descent (IBD) segments calling. (A) Spatial simulation framework in SLiM to produce tree sequences (Haller & Messer, 2023). Individuals are distributed on a two-dimensional habitat map defining suitable areas. Local population density is regulated through a local carrying capacity parameter (*K*) and density-dependent interactions acting at a spatial scale (σ_X_). Reproduction is governed by spatially constrained mate choice, with interaction strength according to a mating scale (σ_M_). Offspring are dispersed from their parents following a probabilistic dispersal kernel characterized by a dispersal distance (σ_D_). (B) Demographic scenarios. Exploration of a range of local carrying-capacity density values and offspring-mother dispersal distance (C) Ecological parameters. 1) Spatial structure is defined either by patchy habitat suitability or by continuous environmental gradients; 2) Offspring dispersal follows alternative dispersal kernels: Student or Laplace distributions; 3) Life history strategies differ in lifespan, age-specific survival and reproduction probabilities and hermaphrodism. (D) Methodological parameters. 1) Sampling effort varies from a small number of individuals to near-exhaustive population sampling; 2) Genealogical information is analysed at different temporal depths. 3) Different ARG inference methods are tested. (E) Inference of IBD segments from simulated genealogies. Tree sequences generated by SLiM are analysed using *tskit* (Kelleher et al., 2016; Wong et al., 2024) to trace shared ancestry between sampled individuals through time. Genealogical relationships are inferred from IBD segments. The reference configuration is defined as average conditions: a homogeneous environment, a carrying capacity K=1, a dispersal variance σ=1, a gaussian dispersal kernel, a sample size of hundred individuals, a temporal depth of two hundred generations.

### 2.1 | Spatial genetic model

We implement a spatial genetic model in *SLiM V5.0* ((Haller & Messer, 2023), Figure 1A) that integrates several key components: (i) an individual-centred genetic model with recombination, (ii) a non-Wright-Fisher spatial population dynamics with density-dependent regulation, (iii) a dispersal kernel defining the spatial scale of dispersal, (iv) a two-dimensional spatial map explicitly defining local habitat availability and dispersal, and (v) a life table describing age-specific survival and fecundity rates.

#### Individual-centred genetic model

Each individual has its genome composed of a single chromosome (L = 10^8^ pb) subject to uniform recombination during meiosis (recombination rate r = 10^-8^ crossover per bp per generation).

#### Spatial map

The habitat is represented as a square grid of dimensions W x W. Each grid cell can be associated with a local value for the carrying-capacity parameter (K) or the dispersal parameter (σ_D_), allowing spatial heterogeneity in local habitat size and spatial scale of dispersal.

#### Spatial population dynamics

Population regulation is density-dependent and spatially explicit. At each reproduction cycle and for each individual, we evaluate the local density of neighbouring individuals (*neighbourhood density)*, defined as the sum of competitive interaction forces calculated from a Gaussian density with standard deviation σ_X_ and a maximum interaction distance of 3σ_X_. The number of offspring per mating event follows a Poisson distribution, whose expectancy is governed by a Beverton-Holt relationship:

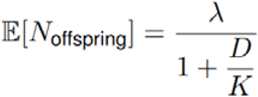

where λ is the maximum fecundity at low density (λ=2), D is the *neighbourhood density,* and *K* represents the local carrying-capacity parameter. This formulation captures local intraspecific competition, with fecundity decreasing asymptotically as density increases. The mate of each individual in each time step is selected randomly, with probability proportional to a Gaussian density with standard deviation σ_M_ and a maximum interaction distance of 3σ_M_. When a mating partner is available, an offspring is produced through sexual reproduction.

#### Spatial dispersal process

Each offspring disperses from the first parent’s location by an independent random displacement in each dimension that is Gaussian distributed with mean zero and standard deviation σ_D_. Offspring dispersing outside the bounds of the habitat are removed from the simulation.

#### Life cycle and age structure

First, we considered a simple non-overlapping discrete-generation model, where all individuals have an equal chance to reproduce and produce offspring before dying. Then, we used an age-structured model where iteroparity and overlapping generations emerge from life-table parameters defining the survival probability and fertility at age x.

### 2.2 | Tree sequence

Simulations output recombinant genealogies in the form of an ancestral recombination graph (*ARG*), encoded as a tree sequence, which provides direct access to the full coalescent history with recombination along the genome (Figure 1D). Each simulation was initiated with 10,000 randomly distributed ancestral individuals and ran for 40,000 generations. Finally, we used a “recapitation” strategy that employs a coalescent simulation in *msprime* V1.0 (Baumdicker et al., 2022) to make ancestor lineage coalesce at all genealogical trees. At the end of each simulation, we randomly subsampled from ten to one thousand individuals and then simplified the tree sequences to retain only nodes ancestral to the sampled genomes.

We extracted Identity-by-Descent (IBD) segments from the tree sequence using a custom script called *ibd-extract-stitch.py* that relies on the *ibd-segments* function of *tskit* V0.6.2 (Kelleher et al., 2016; Wong et al., 2024) but conserves the continuity of IBD segments distributed across multiple adjacent trees. The function returns a data frame of all the pairs of IBD segments shared by all the pairs of individuals, alongside their chromosomal coordinates and age of the most recent common ancestor (in generations). This function was applied with the *max_time* parameter to retain only IBD segments inherited from recent common ancestors that occurred below a given time threshold.

### 2.3 | Demographic inference from IBD segments

We performed demographic inference from the resulting IBD segments using two complementary methods: *IBD-Analysis* (Ringbauer et al., 2017) and *MAPS* (Al-Asadi et al., 2019). Both approaches rely on the principle that the probability of sharing an IBD segment of a given length between two individuals depends on the time to their most recent common ancestor and the geographic distance between them resulting from multigenerational dispersal. These relationships reflect recent demographic connectivity and are jointly shaped by population density and dispersal processes. *IBD-Analysis* provides a single estimate of these demographic parameters across the entire landscape, whereas *MAPS* infers spatial variation in these parameters and outputs dispersal and density maps.

#### 2.3.1 | IBD-Analysis

We first used *IBD-Analysis* to infer density and dispersal parameters. *IBD-Analysis* fits an isolation-by-distance model describing the expected sharing of IBD segments as a function of pairwise geographic distance and segment age. From this model, two key demographic parameters are jointly estimated: (i) the local effective population density *D*, defined as the density of breeding individuals per unit area contributing to recent coalescent events, and (ii) the effective dispersal σ_e_, which characterises the typical spatial displacement between parent and offspring. To enable a direct comparison between simulated and inferred effective densities, we quantified the effective neighborhood-size obtained in the simulations. We calculated the *neighborhood size* (*N_s_*) as the number of individuals within a circular area of radius 2·σ around each focal individual. Then, we converted the *neighborhood size* into an effective population density *D_e_*using the classical relationship 4πD ^2^, from which the realized effective density in the simulations was obtained as:

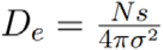

In addition, to enable direct comparison between simulated and inferred dispersal rates, we quantified the effective dispersal rate in our simulations as the root mean squared parent–offspring directional displacement, σ_e_. For each parent–offspring pair, we computed the squared Euclidean displacement d^2^=(x_o_−x_p_)^2^+(y_o_−y_p_)^2^ and estimated:

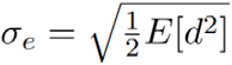

where the expectation is taken over all parent–offspring dispersal events, considering only offspring alive in the final generation. Inference was performed using IBD segments after filtering within a specified length range (2–100 cM), ensuring sensitivity to recent coalescent events and contemporary demographic processes.

#### 2.3.2 | MAPS

Then, we used *MAPS* to estimate spatial variation in dispersal rates and population densities. *MAPS* implements a coalescent-based Bayesian framework that infers local migration rates and population sizes from pairwise sharing of IBD segments summarized in genetic similarity matrices across predefined segment-length bins. We considered IBD segments with lengths ranging from 2 to 100cM. The study area was discretized into a spatial grid composed of 100 demes, which defined the spatial structure used for inference. For each simulation scenario, the MCMC chain was run for 3e6 iterations, with the first 1.5e6 iterations discarded as burn-in to ensure convergence to the equilibrium. We recorded parameter values every 1500 iterations. Convergence across independent chains was also assessed using replicate runs in some scenarios. Inference outcomes were visualised using functions from the *plotmaps* R-package (https://github.com/halasadi/plotmaps), which provides spatial projections of estimated migration and density surfaces.

### 2.4 | Accuracy of demographic inference

We first assessed the accuracy of demographic inferences under a reference configuration characterized by the following simulation parameters: a homogeneous spatial structure with a constant carrying capacity (*K*=1) and variance in dispersal distance (σ=1), a Gaussian dispersal kernel, a sample size of a hundred individuals and a temporal depth of two hundred generations for extracting IBD segments.

#### Variation in carrying capacity (K) or dispersal (σ)

Based on this reference configuration, we explored a range of dispersal and carrying capacity values to assess the robustness and accuracy of demographic inference under varying demographic conditions (Figure 1B). We examined both parameters’ variations across low (0.5), intermediate (2, 3) and high (4, 5) values.

### 2.5 | Robustness of inference to ecological parameters

We explored the impact of ecological parameters to evaluate the robustness of demographic inferences to both demographic and biological complexity (Figure 1C). We examined three main factors: (i) spatial heterogeneity in local population density or dispersal variance, (ii) the shape of the dispersal kernel, and (iii) life cycle complexity. Together, these factors provide a more realistic approximation of biological species evolving in natural environments.

#### Habitat heterogeneity

We modified habitat spatial structure by adjusting the spatial distribution of local carrying-capacity or dispersal according to the pixel values of the environmental map, scaled between zero and one. High pixel values corresponded to higher local values of *K* or σ, and inversely. We simulated two types of spatial heterogeneity – gradient heterogeneity and patchy heterogeneity – to assess how environmental structure influences the accuracy of demographic inference.

#### Dispersal kernel

We evaluated alternative dispersal kernels, including *Laplace* and *Student’s t* distributions. These distributions, characterised by heavier tails than the Gaussian distribution, allow for more frequent long-distance dispersal events and were used to assess the robustness of demographic inference to deviations from the assumed normal distribution of dispersal distances.

#### Life history complexity

We introduced life-history complexity by explicitly incorporating age structure and sequential hermaphrodism into the simulations. We included age-specific survival and fecundity, representing intermediate and long-lived marine fish life history strategies. For each strategy, we combined age structure with sequential hermaphroditism characteristic of focal species: protandry for the intermediate-lived strategy, approximating the life cycle of *sparids* (*Diplodus sargus*), and protogyny for the long-lived strategy, approximating the life cycle of *epinephelids* (*Epinephelus marginatus*). These scenarios allowed us to evaluate the joint effects of age structure and reproductive mode on the accuracy of demographic inference.

### 2.6 | Robustness of inference to methodological parameters

Additionally, we investigated the impact of methodological parameters to evaluate the robustness of demographic inference to analytical choices (Figure 1D). We considered three main factors: (i) sampling size, (ii) temporal depth of IBD segments and (iii) ARG inference method. These parameters directly shape the quantity, quality and structure of information contained in the IBD segments and therefore influence the ability of the methods to accurately estimate the effective population density *D_e_*and the dispersal parameter σ.

#### Sample size

We used three contrasting sampling schemes to evaluate the impact of sample size (*S*) across two orders of magnitude: from a small sample (*S*=10 individuals) to a large sample size (*S*=1000 individuals), in order to quantify how the amount of genetic information available affects the robustness of demographic inference.

#### Temporal depth

We examined three temporal depth thresholds (*t*=50, 100 and 400 generations) to assess their effects on the estimation of local population density and effective dispersal.

#### ARG inference method

We compared two ancestral recombination graph (ARG) inference methods – *tsinfer* coupled with *tsdate* (Kelleher et al., 2019b; Pope et al., 2026; Wohns et al., 2022) and *Singer* (Deng et al., 2024) – to evaluate how differences in genealogical reconstruction affect demographic estimates compared with the ground true ARG. More specifically, our objective was to assess whether ARG-based approaches can provide a reliable alternative to standard IBD-calling pipelines for downstream demographic inference. Because IBD segment detection depends on the underlying inferred ARG, variation among methods can influence the number, age and length of detected segments and thus the estimation of effective population density (D) and effective dispersal (σ_e_). We were unable to include *Relate* (Speidel et al., 2019) in this comparison because it assigns distinct node identifiers to each marginal tree along the genome, preventing our IBD stitching function from merging IBD segments across adjacent trees. Specific methodological development would therefore be required to make *Relate* outputs compatible with our framework.

### 2.7 | Evaluation of inference accuracy and robustness

We evaluated the accuracy of both methods on simulated datasets, using the *root-mean-square error* (RMSE) which quantifies the average deviation between the inferred parameter values to their corresponding true values used in the simulation. For each replicate, RMSE was computed as the root of the mean squared difference between estimated and true values:

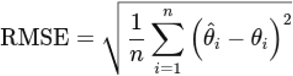

where θ_i_^^^ denotes the estimated parameter value for replicate i and θ the corresponding true value. RMSE values were computed separately for each simulation scenario, parameter, inference method and replicate. Lower RMSE values indicate higher inference accuracy, reflecting closer agreement between inferred and simulated demographic parameters.

To quantify the relative performance of each simulation scenario, we compared results obtained under each tested configuration with those from the reference configuration (calculated as the mean RMSE across replicates for each simulation scenario, inference method and parameter). For this, we used the relative difference, which captures how inference accuracy deviates from the baseline scenario across simulation conditions, parameters and inference methods. A positive value indicates higher estimation error relatively to the reference configuration, and conversely a negative value indicates improved performance relative to the reference configuration.

### 2.8 | Evaluation of spatial pattern inferred by MAPS

We assessed *MAPS* performance by comparing estimated spatial surfaces to the simulated parameter values (effective density D_e_ and dispersal distance σ_e_) for each condition and replicate. This framework allowed us to evaluate whether *MAPS* introduces artificial spatial structure in its demographic parameter estimates.

Under homogeneous landscape, where parameter values are spatially constant by design, any spatial structure detected in *MAPS* estimates necessarily reflects methodological artefacts rather than biological signal. We quantified this artefactual spatial structure using Moran’s *I* statistic (Moran, 2026), computed separately for effective D_e_ and σ_e_ estimates. In addition, spatial confounding between parameters was assessed using a bivariate Moran’s *I* calculated from residuals of D_e_ and σ_e_ (defined as the difference between *MAPS* estimates and the corresponding true simulated values). Positive values indicate that estimation errors for both parameters are spatially co-localized.

For heterogeneous landscapes, we evaluated *MAPS* ability to recover the pattern of the simulated spatial structure. This was quantified using a bivariate Moran’s *I* statistic (Wartenberg, 1985) computed between simulated and estimated parameter surfaces for D_e_ and σ_e_ separately. High positive values indicate that areas of high and low parameter values are spatially matched between simulated and inferred surfaces, whereas lower values indicate weaker reconstruction of the underlying spatial pattern. To assess the influence of background areas with no available habitat or impossible dispersal, analyses were performed both on the complete landscape and after masking cells that cannot support individuals (zero carrying capacity K or no possible dispersal).

## 6 | RESULTS

For each simulation replicate (tens replicate per condition), the output consisted of a dataset of pairwise IBD segments that was subsequently used for demographic inference. The reference simulation (homogeneous landscape with K=1, σ=1, Gaussian dispersal kernel, sample size of hundred individuals and temporal depth of two hundred generations) yielded an average of 31.420 IBD segments. Across all simulated conditions, the number of detected IBD segments ranged from 269 to 3.179.129, reflecting the strong effects of all tested parameters on the amount of detectable IBD segments. As expected, the distributions of IBD segment lengths and ages were consistent with theoretical predictions, with longer IBD segments corresponding to more recent common ancestors (Figure S1).

### 3.1 | *IBD-Analysis* results

#### 3.1.1 | Evaluation of inference accuracy

Overall, *IBD-Analysis* provided relatively precise estimates of demographic parameters across the tested range of values under homogeneous landscapes. Simulated and estimated values of effective density (*D*_e_) showed a strong linear relationship (Figure 2A; slope=0.956, intercept=-0.301, p<0.001), indicating close agreement between inferred and true values. However, the negative intercept indicates a slight systematic underestimation, and precision decreased at higher values of *D*_e_ due to increased variance across replicates. Similarly, estimates of effective dispersal variance (σ*_e_*) were strongly correlated with simulated values (Figure 2B; slope=0.926, intercept=-0.260, p<0.001), although the negative intercept again indicates a slight systematic underestimation. Extending the range of dispersal values revealed the emergence of a plateau at high dispersal levels (σ*_e_* > 10), consistent with a regime where dispersal becomes increasingly difficult to estimate when gene flow is sufficiently high that spatial genetic structure is eroded over multiple generations (Figure S2).

**Figure 2.**
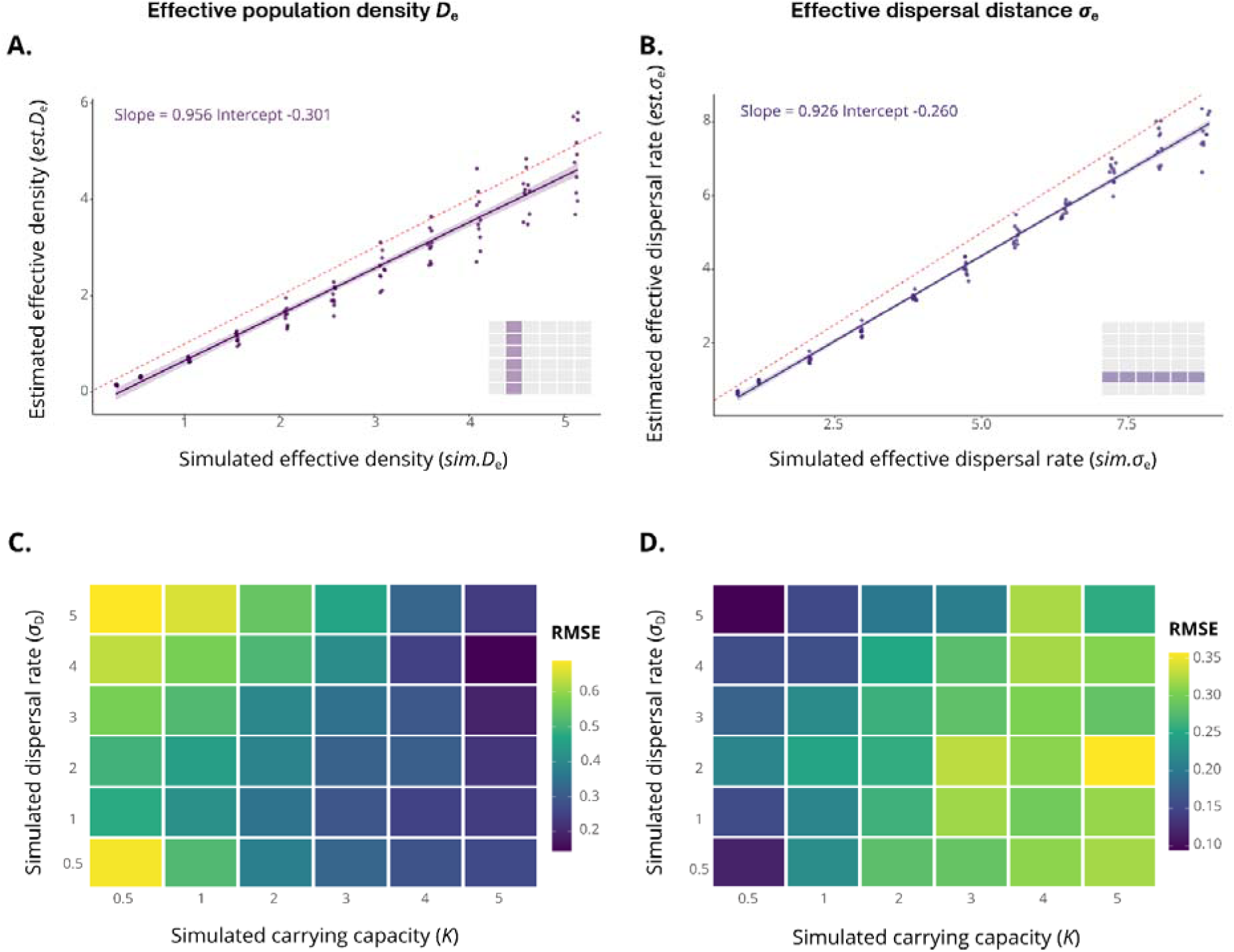
– Accuracy of demographic parameter inference using *IBD-Analysis*. Left panels: Effective population density (*D*_e_). Right panels: Effective dispersal distance (σ_e_). Effective density *D*_e_ depends on the simulated carrying capacity (*K*) and effective dispersal σ_e_ depends on the simulated dispersal distance (σ*_D_*), relating the simulation parameters to their corresponding effective demographic parameters. **(A-B)** Relationship between simulated parameter values and their estimates inferred from IBD segments using *IBD-Analysis*. A dot represents one replicate; solid lines correspond to linear regressions between simulated and estimated values, with shaded areas indicating 95% confidence intervals. The dashed red line represents the 1:1 relationship. **(C-D)** Relative root mean squared error (RMSE) of parameter estimates across combinations of simulated carrying capacity (*K*) and dispersal distance (σ*_D_*). Heatmaps illustrate how inference accuracy varies jointly with demographic conditions.

The RMSE patterns across combinations of *D*_e_ and σ*_e_* revealed contrasting sensitivities for the two parameters. Effective density estimates were generally less accurate at low *D*_e_ values, indicating that errors represent a larger relative deviation from the true value in low-density populations (Figure 2C). Conversely, RMSE for effective dispersal increased with increasing D_e_, indicating that higher effective densities make accurate estimation of σ*_e_* more challenging, although this effect was weaker than the sensitivity observed for D_e_ estimates (Figure 2D).

#### 3.1.2 | Evaluation of robustness to methodological and ecological parameters

##### Impact of ecological parameters

Ecological parameters had strong and heterogeneous effects on inference accuracy. For effective density (Figure 3A), several ecological scenarios resulted in increased signed RMSE relative to the reference configuration, indicating systematic deviations from the true values. The strongest effects were associated with life-history characteristics, particularly under intermediate and long-life cycles, where increasing maximum fecundity further reduced the inference accuracy, especially for the intermediate lifespan protandrous life-cycle scenarios. In comparison, spatial heterogeneity across the landscape had a lower impact on effective density estimates and was mostly associated with heterogeneous density with little impact of heterogeneous dispersal.

**Figure 3.**
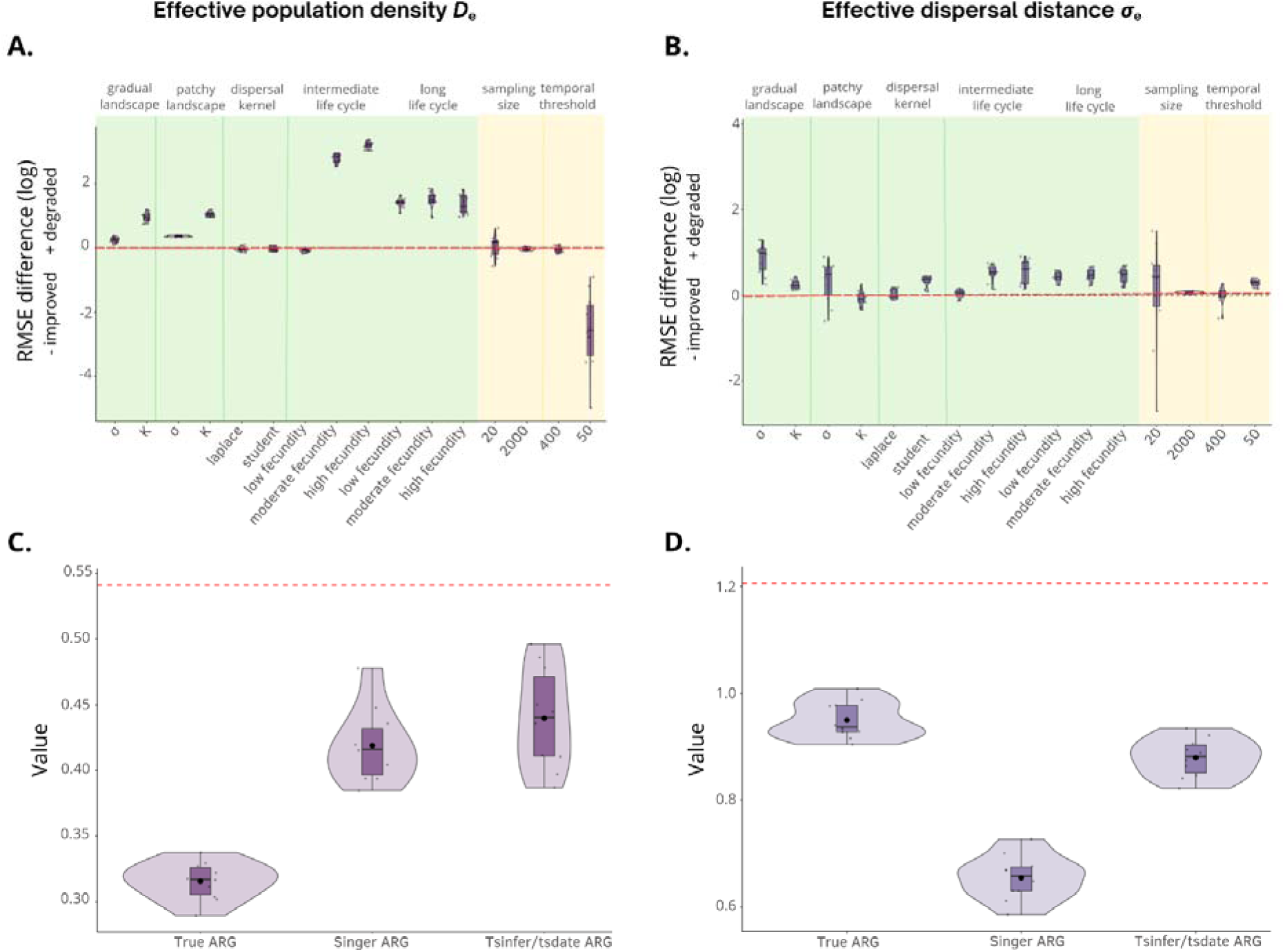
– Robustness of parameter inference across tested conditions using *IBD-Analysis*. Lefts panels: Effective population density (*D*_e_). Right panels: Effective dispersal distance (σ_e_). **(A-B)** Log-transformed relative root mean squared error (RMSE) of parameter estimates under each tested configuration compared with the reference configuration. Relative RMSE was calculated as the difference between the RMSE obtained under a given condition and that obtained under the reference configuration. The reference configuration is defined as average conditions: a homogeneous environment, a carrying capacity *K*=1, a dispersal variance σ=1, a gaussian dispersal kernel, a sample size of hundred individuals, a temporal depth of two hundred generations. The green area represents the ecological parameters, and the yellow area represents the methodological parameters. Positive value indicates a degradation of demographic inference relative to the reference condition. The red dashed line denotes no change relative to the reference configuration. **(C-D)** Value of parameters estimates in the reference scenario obtained following IBD-segment detection based on the true ARG, Singer ARG and tsinfer/tsdate ARG. The red dashed line represents the true parameter value.

For effective dispersal (Figure 3B), ecological parameters also significantly influenced inference accuracy. In contrast to density estimates, spatial heterogeneity in dispersal across the landscape was the main factor affecting dispersal estimates, whereas life-history characteristics had comparatively weaker effects. The effect of the dispersal kernel was only detected using the *Student* distribution with the heavier tail, which led to reduced inference accuracy compared with the reference scenario.

##### Impact of methodological parameters

Methodological parameters had generally weaker effects on inference performance than ecological parameters. Variation in sample size (*S*) and the temporal threshold used for IBD segment selection (*t*) had relatively limited influence on estimates of both effective density and dispersal (Figure 3A, B). Among these factors, reduced sampling size had the strongest effect, primarily for dispersal estimates where diploid sample sizes of 20 chromosomes resulted in increased variability among estimates (Figure 3B). Interestingly, reducing the time threshold for IBD segment selection increased the relative precision of density estimates, but resulted in a smaller decrease in dispersal estimation accuracy.

Using IBD segments extracted from reconstructed ARGs introduced an additional source of uncertainty, even though ARGs were inferred from ideal genome data without genotyping or phasing errors. Parameter estimates obtained from different ARG reconstruction methods consistently deviated from those obtained using the true ARG. Comparisons between inferences based on reconstructed and true ARGs revealed a general tendency toward lower demographic parameter estimates, but with different magnitudes depending on parameters (Figure 3C-D). For D_e_, estimates based on *Singer*-reconstructed ARGs were closer to those obtained from the true ARG than those based on *tsinfer/tsdate* ARGs. Conversely, for σ_e_, estimates derived from *tsinfer/tsdate* were closer to the true ARG estimates than those obtained from *Singer*, possibly because biases in node age and IBD segments length partially compensated for errors introduced during ARG reconstruction. Overall, ARG reconstruction introduced measurable deviations in demographic parameter estimates relative to the true ARG, highlighting the influence of genealogical reconstruction uncertainty on the robustness of IBD-based demographic inference.

Overall, ecological complexity had a stronger influence on inference accuracy than methodological parameters, with life-history variation and density heterogeneity primarily affecting estimates of D, while dispersal heterogeneity primarily affected estimates of σ.

### 3.2 | *MAPS* results

Overall, *MAPS* recovered broad variation in demographic parameters across homogeneous landscape simulations, but inference performance differed markedly between effective density (D_e_) and effective dispersal (σ_e_). Estimates of D_e_ showed a relatively strong linear relationship with simulated values, however, regression slopes consistently indicated increasing overestimation at higher density values (Figure S3A; slope=2.53, intercept=-0.09). In contrast, estimates of σ_e_ were more overestimated but also showed substantially higher variability across replicates (Figure2B; slope=10.31, intercept=2.82), resulting in lower precision, especially at moderate to high dispersal values. RMSE patterns further revealed distinct sensitivities between parameters: error in D_e_ estimates were highest under low carrying capacity (K), whereas error in σ_e_ increased markedly with increasing dispersal rate, especially at higher values of *K* (Figure S3C-D). Sensitivity analyses indicated that, as observed for *IBD-Analysis*, ecological parameters had a stronger influence on inference performance than methodological parameters. Spatial heterogeneity and complex life-history scenarios produced the largest deviations from the reference configuration (Figure S4A-B). In addition, ARG reconstruction affected parameter inference, with both reconstruction methods leading to overestimation of D_e_ but underestimation of σ relative to estimates obtained from the true ARG (Figure S4C-D).

#### 3.2.1 | Spurious spatial structure in homogeneous landscapes

Despite the absence of true spatial variation in demographic parameters, *MAPS* produced strongly spatially autocorrelated parameter surfaces under homogeneous landscape conditions (Figure 4). Moran’s *I* values were consistently high for both effective density (D_e_; Figure 4A) and effective dispersal (σ_e_; Figure 4B), generally exceeding 0.9 across combinations of carrying capacity (K) and dispersal rate (σ_D_). This spatial pattern was magnified under combinations of low K and high σ_D_, or high K and low σ_D_, where Moran’s *I* values approached one, and slightly reduced under high K and high σ_D_.

**Figure 4.**
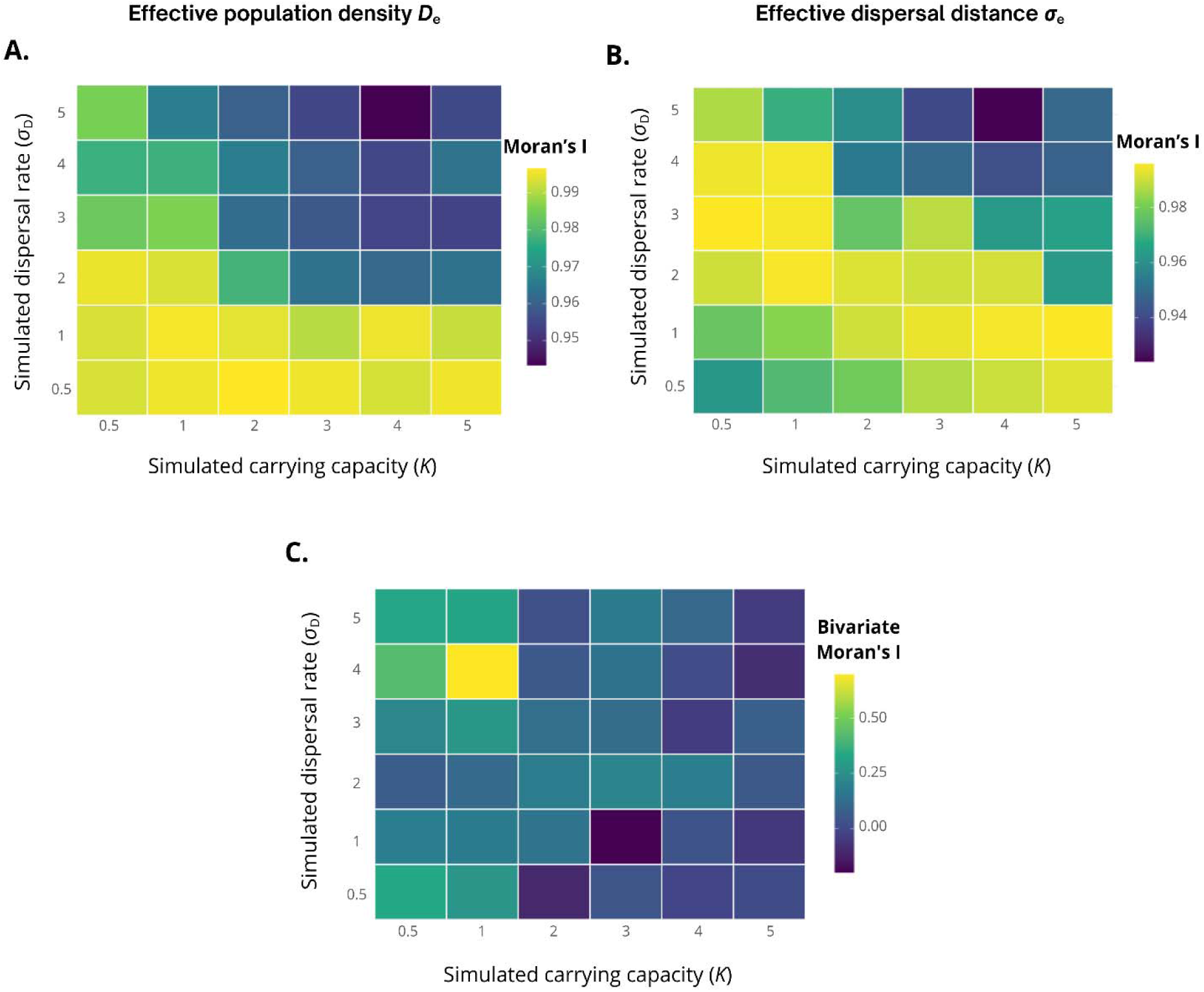
– Spatial autocorrelation of *MAPS* parameter estimates under homogeneous landscape conditions. A: Effective population density (*D*_e_). B: Effective dispersal distance (σ_e_). C: Cross-parameter spatial confounding. Effective density *D*_e_ depends on the simulated carrying capacity (*K*) and effective dispersal σ_e_ depends on the simulated dispersal distance (σ*_D_*), relating the simulation parameters to their corresponding effective demographic parameters. **(A-B)** Moran’s *I* statistic computed on *MAPS* estimates of *D*_e_ and σ_e_ respectively, across combinations of simulated carrying capacity (*K*) and dispersal rate (σ_D_). **(C)** Bivariate Moran’s *I* statistic computed between the residuals of *D*_e_ and σ_e_ estimates across spatial positions. Positive values indicate that *MAPS* commits spatially co-localized estimation errors for both parameters simultaneously. bivariate Moran’s *I* statistic computed between the residuals of D_e_ and σ_e_ estimates across spatial positions. Positive values indicate that *MAPS* commits spatially co-localized estimation errors for both parameters simultaneously. Each cell represents the median value across replicates for the corresponding parameter combination.

The bivariate Moran’s *I* calculated from residuals of D_e_ and σ_e_ estimates revealed that spatial confounding between the two parameters depended strongly on simulation conditions (Figure 4C). Values ranged from near zero to approximately 0.6, indicating substantial variation in the extent to which estimation errors were spatially co-localized. Residual correlations were strongest at intermediate combinations of K and σ_D_, where *MAPS* tended to overestimate or underestimate both parameters in the same regions. Conversely, weak or negative values were observed under very low dispersal or high carrying capacity, suggesting limited spatial association, or even spatial opposition, between estimation errors in D_e_ and σ_e_.

#### 3.2.2 | Landscape-dependent recovery of spatial structure

Under heterogeneous landscape conditions, *MAPS* showed substantial variation among simulation conditions and replicates in its ability to recover simulated spatial structure (Figure 5). While some simulations replicate successfully captured the expected spatial patterns, yielding relatively high positive bivariate Moran’s *I* values (approximately 0.8 for D_e_ and 0.5 for σ_e_), others produced values close to zero or even negative, indicating weak or no correspondence between the inferred and simulated landscapes. This variation among replicate simulations was observed across both landscape configurations (gradual and patchy) and demographic parameters (D_e_ and σ_e_).

**Figure 5.**
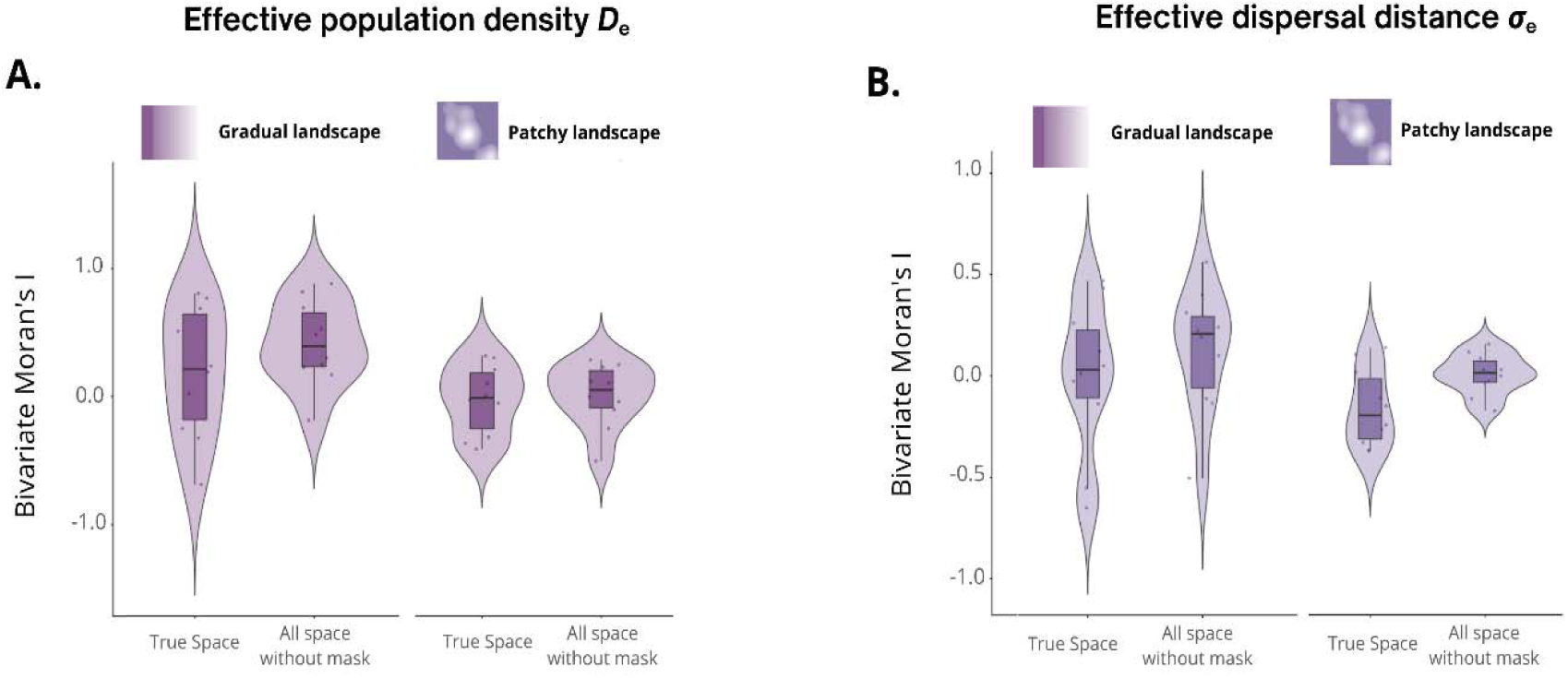
– Spatial autocorrelation of *MAPS* parameter estimates under heterogeneous landscape conditions. **(A)** Bivariate Moran’s *I* statistic computed between simulated and *MAPS*-estimated *D*_e_ surfaces across combinations of heterogeneous landscape types (patchy and gradient) comparing analyses performed on the full simulated space *versus* the “true” space obtained without masking cells that cannot support individuals. **(B)** Bivariate Moran’s *I* statistic computed between simulated and *MAPS*-estimated σ_e_ surfaces across the same conditions. Positive values indicate spatial agreement between simulated and inferred surfaces.

For effective density (Figure 5A), bivariate Moran’*I* values were generally positive, suggesting that *MAPS* recovered part of the simulated spatial structure in several simulations. Analyses performed over the complete spatial domain without masking unsuitable areas tended to produce slightly higher Moran’s *I* values than analyses restricted to biologically available habitat (“true space”), although this difference remained modest.

A similar pattern was observed for effective dispersal (Figure 5B), although spatial correspondence was generally weaker than for effective density. Estimates obtained from the complete spatial domain tended to produce Moran’s *I* values closer to zero or slightly positive, whereas analyses restricted to true space more frequently resulted in negative values, particularly in patchy landscapes. However, these tendencies remained highly variable among simulation replicates, with substantial overlap between distributions.

Generally, gradual landscapes tended to produce higher bivariate Moran’s *I* values than patchy landscapes, irrespective of whether analyses were conducted in true space or over the complete spatial domain. However, substantial overlap among distributions indicates that this tendency was weak relative to the large variation among simulation replicates.

Overall, as observed for *IBD-Analysis*, ecological complexity had a stronger influence on *MAPS* inference accuracy than methodological parameters. *MAPS* recovered broad demographic variation, but effective density (D_e_) was estimated more reliably than effective dispersal (σ_e_). Moreover, spatial inference remained sensitive to landscape structure, generating spurious spatial patterns in homogeneous landscapes and only partially recovering simulated spatial variation under heterogeneous landscapes.

## 7 | DISCUSSION

Estimating demographic parameters such as effective population density and dispersal is central to conservation biology yet remains challenging because their effects on spatial genetic structure are often difficult to disentangle. By exploiting the genealogical information contained in identical-by-descent (IBD) segments, recent approaches offer new opportunities to disentangle the respective contributions of density and dispersal to spatial genetic structure. Here, we evaluated and compared two approaches, *IBD-Analysis* (Ringbauer et al., 2017) and *MAPS* (Al-Asadi et al., 2019), across a broad range of simulated spatial scenarios. Our results revealed a trade-off between robustness and spatial resolution: *IBD-Analysis* provided robust global estimates of effective density and dispersal, whereas *MAPS* sacrificed some accuracy in global parameter estimation, particularly for dispersal, to recover local variation in demographic processes. This trade-off becomes especially evident in heterogeneous landscapes, where spatial variation provides the ecological signal that enables local inference, while simultaneously increasing complexity that limits model performance. Rather than competing alternatives, the two approaches provided complementary perspectives on spatial population structure. Together, these findings help guide the choice of IBD-based inference methods depending on the ecological context and highlight their complementary potential for investigating recent demographic connectivity and informing conservation.

### Complementary views of space: global robustness versus local resolution

Although both methods rely on the same genealogical information, they differ fundamentally in how they represent space. *IBD-Analysis* integrates genetic information across the landscape to estimate global demographic parameters averaged over the study area, favoring robustness and statistical power. MAPS, in contrast, explicitly models spatial variation and aims to reconstruct local heterogeneity in demographic processes, providing greater spatial resolution but increased sensitivity to spatial complexity. This difference reflects a broader conceptual distinction in spatial population genetics between global estimators, which prioritize robustness and statistical power, and spatially explicit approaches, which favour local resolution and ecological interpretability (Bradburd & Ralph, 2019). Rather than competing frameworks, these approaches address different inferential objectives and should be viewed as complementary depending on the ecological context and the spatial scale of the biological question.

### Overall shared performance of IBD-based approaches

Both methods estimated effective population density across a wide range of demographic conditions, with *IBD-Analysis* consistently showing lower bias than MAPS. This indicates that IBD segments carry strong and robust information for recovering variation in effective density. In contrast, effective dispersal was harder to estimate. *IBD-Analysis* remained accurate across a broad range of dispersal values (0.5-15% of range dimensions) but reached a plateau at high dispersal values (≥20% of range dimensions), whereas *MAPS* showed lower accuracy and greater variance, particularly under high dispersal conditions. This likely reflects the progressive loss of local spatial signal in highly connected populations, where related individuals are spread over larger geographic distances. Differences in performance became more pronounced when spatial heterogeneity was introduced, indicating that landscape structure is a major determinant of inference accuracy.

### Spatial heterogeneity as the main determinant of performance

Spatial heterogeneity represents both the ecological signal that spatially explicit approaches aim to recover and the main challenge that limits inference accuracy. While both methods performed similarly under homogeneous conditions, their behaviour diverged strongly once demographic parameters varied across space, revealing a fundamental trade-off between spatial resolution and robustness.

For *IBD-Analysis*, spatial heterogeneity acts primarily as a source of systematic bias. Because the method integrates information across the entire landscape, local variation in density or dispersal is summarized into a single landscape-wide estimate. This averaging process provides robustness by reducing sensitivity to local stochastic variation, but it also limits the ability to capture spatial extremes or fine-scale demographic structure. As a result, heterogeneous landscapes lead to deviations between estimated and true values, as local variation is incorporated into a global summary. This behaviour mirrors the loss of demographic information that occurs when effective population size is estimated from genetically structured populations analyzed as a single panmictic unit (Mazet et al., 2016; Waples, 2016; Waples & Gaggiotti, 2006). Although *IBD-Analysis* includes the *MLE_barrier* function to account for spatial barriers, this approach requires prior knowledge of barrier location. Such information is rarely available in natural systems, particularly in marine environments where dispersal barriers are often cryptic and difficult to identify *a priori* (Benestan et al., 2021; Cowen & Sponaugle, 2009; Palumbi, 2003).

In contrast, *MAPS* explicitly attempts to reconstruct spatial variation and therefore provides spatial resolution, but at the cost of increased sensitivity to landscape complexity. Its performance varied markedly among simulation replicates: some successfully recovered the simulated spatial patterns, whereas others showed weak to no correspondence, or even inverse relationships with the true landscape. This variability suggests that datasets simulated under identical demographic scenarios may contain different amounts of exploitable spatial information, influencing the ability of *MAPS* to distinguish signal from stochastic variation.

Landscape configuration further shaped *MAPS* performance. Gradient landscapes, characterized by continuous variation in demographic parameters, were more reliably reconstructed than patchy landscapes, where abrupt discontinuities generated locally inconsistent patterns that are difficult to represent within a spatially continuous framework. Across scenarios, effective dispersal (σ_e_) was consistently reconstructed less accurately than effective density (D_e_), reflecting the greater difficulty of estimating dispersal from spatial patterns of IBD sharing (Rousset, 1997).

In homogeneous landscapes, *MAPS* showed an additional limitation: in the absence of true spatial variation, stochastic variation in IBD sharing was interpreted as a spatial structure, generating artificial spatial autocorrelation. This reflects an intrinsic property of spatially explicit models, which are designed to recover spatial variation and may therefore overfit noise when spatial structure is absent.

Beyond landscape heterogeneity, a more fundamental limitation arises from the shared information content used to infer effective density (D_e_) and effective dispersal (σ_e_). Both parameters shape the same patterns of IBD sharing, making them intrinsically difficult to estimate independently. This problem becomes particularly acute under high dispersal and weak spatial genetic structure, where geographic distance is only weakly associated with genetic relatedness. Under these conditions, the information available in IBD segments becomes insufficient to clearly distinguish the effects of density from those of dispersal.

*MAPS* then tends to compensate uncertainty in one parameter by adjusting the other, resulting in mirrored biases in the inferred spatial surfaces. Such behaviour reflects a fundamental identifiability issue rather than a methodological artefact: when two demographic processes leave similar signatures in the same genealogical signal, uncertainty in one parameter inevitably tends to propagate to the other. Improving the independent estimation of density and dispersal will therefore likely require incorporating additional sources of information, such as ecological data to constrain the model.

Contrary to expectation, restricting the analysis using prior knowledge of biologically accessible habitat (“true space”) did not improve reconstruction accuracy. Instead, analyses performed over the complete spatial domain generally showed slightly stronger correspondence with the simulated landscapes for both effective density (D_e_) and dispersal (σ_e_), although these differences remained modest and highly variable among simulation replicates.

One explanation is that MAPS benefits from estimating demographic surfaces over a continuous spatial domain, where a more regular lattice provides improved spatial regularization and stability of parameter estimates. Restricting the analysis to occupied habitat, in contrast, adds boundaries that may increase edge effects and reduce the spatial information available for smoothing parameter surfaces. Whether this behaviour reflects a general property of *MAPS* or is specific to the landscapes considered here remains to be determined.

Overall, spatial heterogeneity emerged as the principal determinant of inference performance. Depending on landscape complexity and the inference framework, IBD-based approaches may provide robust global estimates, recover local spatial variation, or generate spurious spatial structure when the underlying signal is weak.

### Uncertainty associated with ARG-based IBD reconstruction

How IBD segments are obtained represents an important methodological consideration for demographic inference. Both *IBD-Analysis* and *MAPS* can be applied to segments detected directly with IBD-calling algorithms (Browning & Browning, 2020; Fournier et al., 2023; Guo et al., 2024; Nait Saada et al., 2020; Seidman et al., 2020; Zhou et al., 2020), but the increasing availability of ancestral recombination graph (ARG) inference methods offers an attractive alternative approach by reconstructing genome-wide genealogies from sequence data (Brandt et al., 2024; Hudson, 1991; Lewanski et al., 2024; Nielsen et al., 2025). Because demographic inference depends on the accuracy of these reconstructed genealogies, we tested whether IBD segments extracted from inferred ARGs provide reliable estimates of recent demographic parameters.

ARG inference remains challenging because sequence variation provides only incomplete information about the underlying genealogy (Deng et al., 2024; Rasmussen et al., 2014; Speidel et al., 2019; Wohns et al., 2022). Errors in reconstructed topologies and node ages distort the inferred distribution of IBD segments by fragmenting or conflating true segments and altering estimates of recent coalescent times (Brandt et al., 2024). These distortions propagate directly into demographic inference because both *IBD-Analysis* and *MAPS* rely on the length distribution of IBD segments. In our simulations, they consistently biased parameter estimates relative to those obtained from the true ARG, leading to overestimation of effective density (*D*_e_) and underestimation of effective dispersal distance (σ_e_). For *D*, this reconstruction-induced bias partially compensated for the slight underestimation observed when *IBD-Analysis* was run using IBD segments extracted from the true ARG, illustrating that different sources of bias can interact in non-intuitive ways.

The magnitude of these biases differed among ARG reconstruction methods, suggesting that method-specific reconstruction errors propagate differently into demographic inference. Previous studies have shown that the algorithm *tsdate* tends to overestimate the age of recent nodes, whereas *Singer* doesn’t (Brandt et al., 2024; Deng et al., 2025). This bias in node-age estimation may partially compensate for its tendency to substantially overestimate IBD length, potentially explaining why *tsinfer/tsdate* algorithms yielded more accurate estimates of dispersal despite larger deviations in other aspects of the reconstructed genealogy compared to *Singer*.

Importantly, our simulations considered an idealized setting in which ARGs were reconstructed from error-free genomic data. In empirical datasets, additional sources of uncertainty such as phasing errors, missing data, genotype calling errors and imputation artefacts, are all expected to further affect ARG reconstruction and, consequently, the detection and inferred properties of IBD segments (Browning & Browning, 2012; Palamara et al., 2012; Ringbauer et al., 2017; Wang et al., 2025).

Overall, ARG reconstruction uncertainty represents an important source of bias for IBD-based demographic inference, with effects that depend on both the demographic parameter being estimated and the reconstruction method used. As ARG-based approaches become increasingly widespread, explicitly accounting for reconstruction uncertainty, and when possible, comparing ARG-derived with directly detected IBD segments, will be essential for robust demographic inference.

### Context-dependent choice of spatial inference methods

The choice between *IBD-Analysis* and *MAPS* should be guided by both the ecological context and the objectives of the study (Figure 6). When demographic processes are expected to be relatively homogeneous across the landscape, *IBD-Analysis* provides the most robust and reliable estimates of effective density and dispersal. Under these conditions, its integrated treatment of space minimizes sensitivity to stochastic spatial variation, whereas *MAPS* may overinterpret local fluctuations in IBD sharing and introduce artificial spatial structure. This scenario is likely to apply to populations inhabiting relatively small or environmentally uniform habitats, where assuming spatial homogeneity is biologically reasonable.

**Figure 6.**
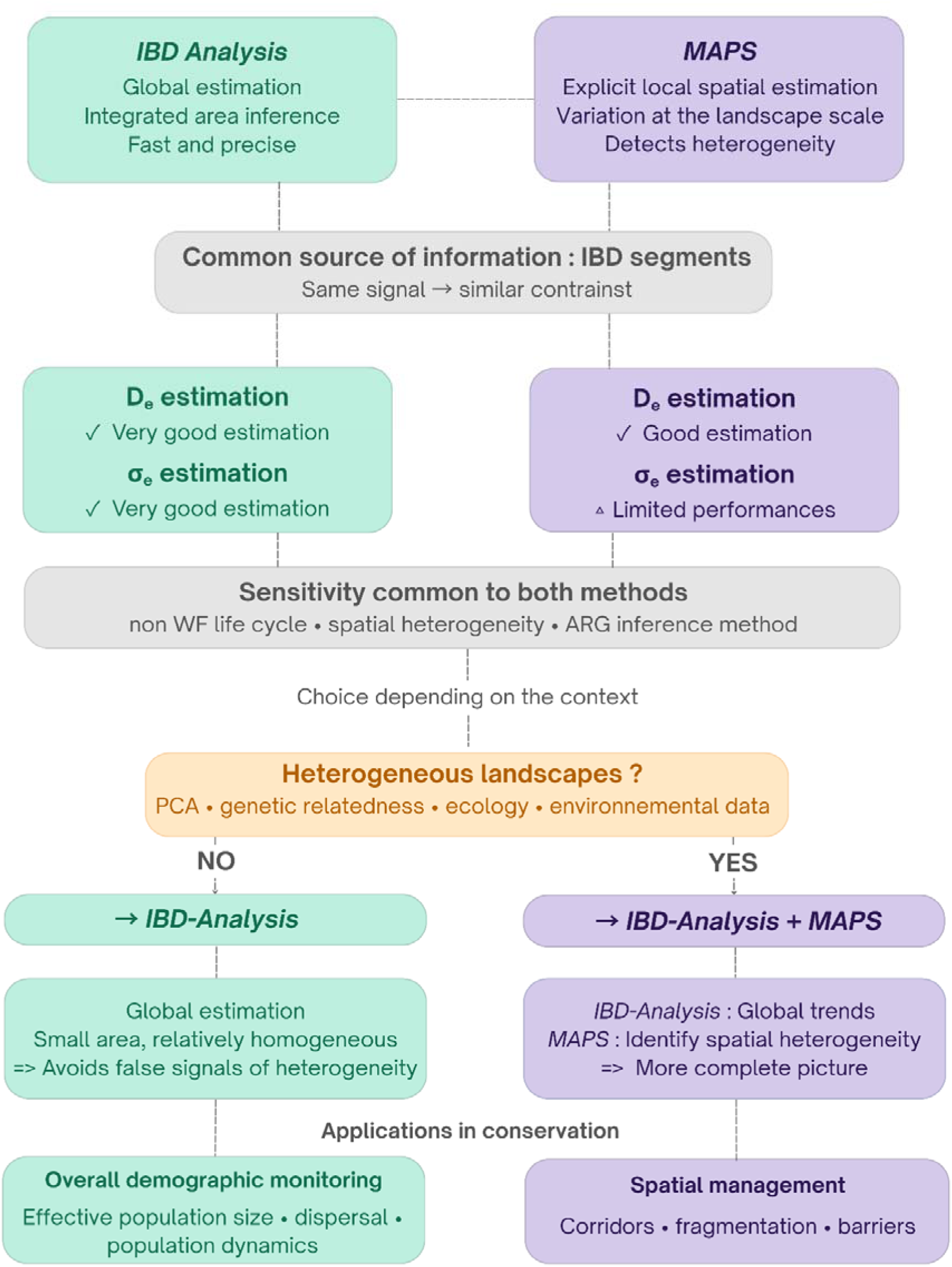
– *IBD-Analysis* vs *MAPS*: A guide to choose the right methodology to estimate demographic parameters.

When demographic processes vary across space, however, the two methods become complementary. *IBD-Analysis* provides a robust summary of average demographic conditions across the study area, whereas *MAPS* can reveal local variation in effective density and dispersal. Although spatial heterogeneity introduces bias into the global estimates produced by *IBD-Analysis* and increases the variability of *MAPS*, combining both approaches may offer a more complete picture of recent demographic connectivity by jointly characterizing global trends and local departures from them.

Prior knowledge of the study system can help guide this choice, based on both genetic and ecological information. Genetic signals, such as spatial structure in relatedness matrices, discontinuities detected by Principal Component Analysis, or isolation-by-distance patterns, may indicate that demographic processes vary across the landscape. Likewise, ecological and environmental knowledge, such as known physical barriers to dispersal, habitat fragmentation, or strong environmental gradients, provides additional support for adopting a spatially explicit framework. In many empirical systems, integrating the complementary information provided by *IBD-Analysis* and *MAPS* is therefore likely to provide the most informative description of recent demographic connectivity.

### From inference to conservation: implications for management

These findings have direct implications for conservation and population management. However, the interpretation of demographic estimates obtained from IBD-based approaches depends on the demographic assumptions underlying the inference models. Departures from Wright–Fisher assumptions, including age structure, sequential hermaphroditism, and high variance in reproductive success, affect the performance of both approaches. Such processes are widespread across taxa and can alter coalescent patterns, weakening the theoretical relationship between IBD sharing and demographic parameters (Charlesworth, 2009; Hudson, 1991). These effects are particularly relevant in marine species, where reproductive success is often highly skewed and effective population size can deviate substantially from census size (Frankham, 1995; Hedgecock & Pudovkin, 2011; Palstra & Ruzzante, 2008; Waples, 2016; Waples et al., 2018). Despite these limitations, IBD-based approaches provide valuable insights into recent demographic processes and can inform conservation strategies when interpreted within an appropriate demographic framework.

When the objective is broad-scale demographic monitoring – such as estimating effective population size or assessing population trends – *IBD-Analysis* provides a robust and interpretable framework. This is particularly relevant for threatened species with small population sizes, where reliable estimates of effective population size are essential for evaluating extinction risk and the strength of genetic drift (Frankham, 1995, 2005; Palstra & Ruzzante, 2008; Waples, 2016).

When the objective is to understand how demographic processes vary across space, MAPS provides complementary spatially explicit information. Beyond estimating average demographic parameters, IBD-based approaches give access to the recent genealogical processes shaping genetic diversity, offering a perspective the spatio-temporal dynamics of the population. This is particularly valuable for local spatial management and conservation, including the identification of barriers to gene flow, ecological corridors and management units (Funk et al., 2012). Traditional spatial genetic methods including isolation-by-distance analyses, barrier detection methods and landscape genetics frameworks (e.g. Currat et al., 2019; Dupanloup et al., 2002; Fitzpatrick et al., 2010; Guillot et al., 2005; Manni et al., 2004; Rousset, 1999), effectively describe patterns of genetic variation (Manel et al., 2003; Manel & Holderegger, 2013) but generally do not disentangle the respective contributions of dispersal and local density. This distinction is fundamental for conservation because similar patterns of genetic connectivity may arise from different demographic processes. Limited connectivity may reflect reduced dispersal, calling for restoring ecological corridors or reducing the impact of barriers, but it may equally result from low population density, requiring demographic reinforcement instead. By disentangling the respective contributions of dispersal and population density, IBD-based approaches provide a powerful framework for linking observed genetic patterns to their underlying demographic processes and, ultimately, for improving conservation and management decisions.

### Conclusion

Our study contributes to the growing field of spatial demographic inference by demonstrating how IBD-based approaches can link genomic variation, genealogical history, and spatial ecological processes. While machine learning approaches provide powerful predictive frameworks, their inference remains constrained by the demographic scenarios represented in their training datasets (Smith et al., 2024). In contrast, IBD-based approaches exploit recent genealogical information and build on earlier methods based on runs of homozygosity (ROHs, (Bertola et al., 2024; Kardos et al., 2017; Ringbauer et al., 2024)), extending the study of recent coalescence from within individuals to shared ancestry among individuals.

Our results demonstrate that IBD-based approaches provide a powerful framework for spatial demographic inference. *IBD-Analysis* offers robust estimates of global demographic parameters, whereas *MAPS* enables the reconstruction of fine-scale spatial heterogeneity by disentangling the contributions density and dispersal, extending the potential of spatial genetic approaches such as EEMS (Bertola et al., 2024; Jones et al., 2021; Marcus et al., 2021; Peter et al., 2020; Petkova et al., 2014; Pimenta et al., 2019) toward a more mechanistic understanding of the processes shaping spatial genetic structure (García-Jiménez et al., 2025). Beyond these current applications, emerging genealogical approaches that infer the spatial location of ancestral individuals and reconstruct historical population dynamics (Deraje et al., 2025; Kelleher et al., 2018; Speidel et al., 2019; Talbot & Bradburd, 2025; Wohns et al., 2022) further highlight the growing potential of these genealogy-based methods for conservation, although improving resolution for recent demographic events remains a major challenge for contemporary population monitoring (Li & Durbin, 2011; Speidel et al., 2019; Terhorst, 2024).

Overall, our findings show that genealogical information provides a powerful bridge between genomic variation and ecological processes. By moving beyond descriptive patterns of genetic structure to infer the demographic mechanisms that generate them. IBD-based approaches open new opportunities for understanding the demographic processes shaping biodiversity across changing landscapes.

## 8 | ACKNOWLEDGEMENTS

The authors thank Harald Ringbauer and Raphaël Leblois for the helpful discussions and feedback. We also thank Romain Pechey and Iago Bonnici for the development of our custom script called *ibd-extract-stitch.py*. We are grateful to the genotoul bioinformatics platform Toulouse Occitanie (Bioinfo Genotoul, https://doi.org/10.15454/1.5572369328961167E12) for providing computing and storage resources.

## 10 | DATA ACCESSIBILITY

All the scripts used to generate the various spatial demographic simulations, obtain the underlying IBD segments and infer the demographic parameters with *IBD-Analysis* and *MAPS* can be found at https://github.com/OceaneEYCHENNE/….

## 11 | AUTHOR CONTRIBUTIONS

- . Eychenne, S. Manel and P.-A. Gagnaire conceived the study. O. Eychenne conducted data analyses and wrote the first draft. All authors contributed to manuscript revisions and approved the final version of the manuscript.

## 12 | CONFLICT OF INTEREST

The authors declare no conflict of interest.

## 14 | FUNDING INFORMATION

This project was supported by the ANR grant DemoMar ANR-24-CE02-4266.

## 14 | ETHICAL STATEMENT

This study is based exclusively on simulated data and did not human participants, animals, or personal data. Therefore, ethical approval and informed consent were not required.

## Notes

### Competing Interest Statement

The authors have declared no competing interest.

