## Supplementary Materials File for "Navigating between global robustness and fine-scale resolution: the potential of demographic inference based on identical-by-descent segments"

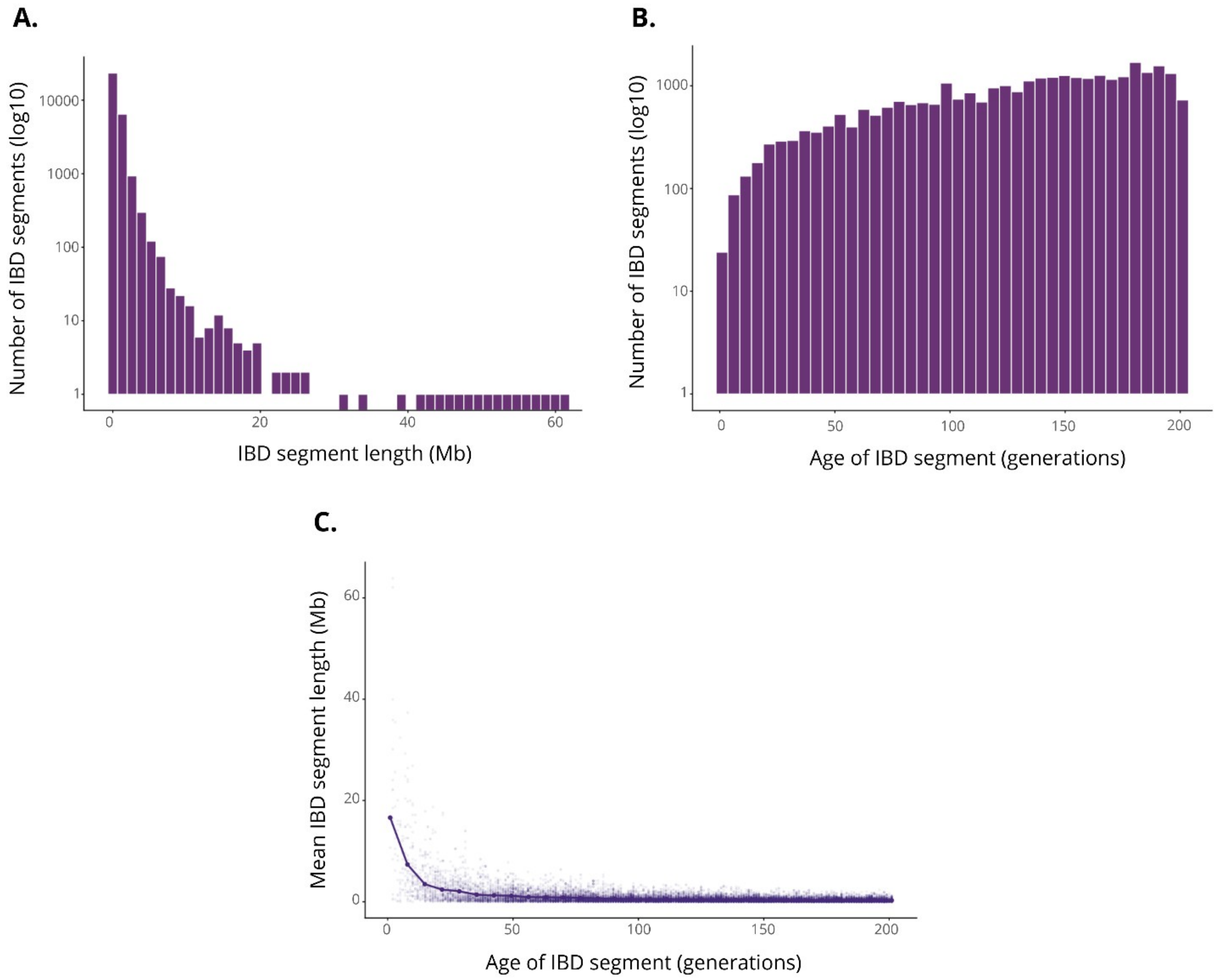


**Figure S1 - Distribution and age of simulated IBD segments. (A)** Histogram of IBD segment lengths bins (Mb), with the y-axis shown on a logarithmic scale. **(B)** Histogram of IBD segment age (generations), with the y-axis shown on a logarithmic scale. **(C)** Relationship between the age of IBD segments (time to the most recent common ancestor, TMRCA, in generations) and their length. Each point represents an individual IBD segment, while the solid line shows the mean segment length computed within age bins.

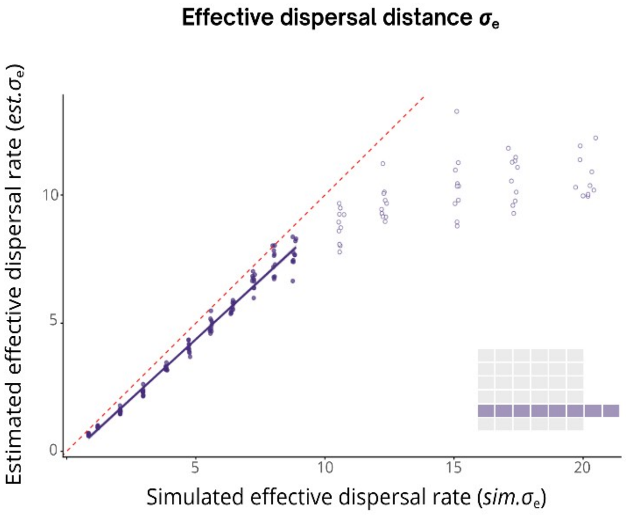


**Figure S2 - Emergence of a plateau in effective dispersal inference using *IBD-Analysis.*** Relationship between simulated parameter values and their estimates inferred from IBD segments using *IBD-Analysis*. A dot represents one replicate; solid lines correspond to linear regressions between simulated and estimated values, with shaded areas indicating 95% confidence intervals. The dashed red line represents the 1:1 relationship. Empty circles represent simulation conditions in which dispersal rate is too high compared to available space.

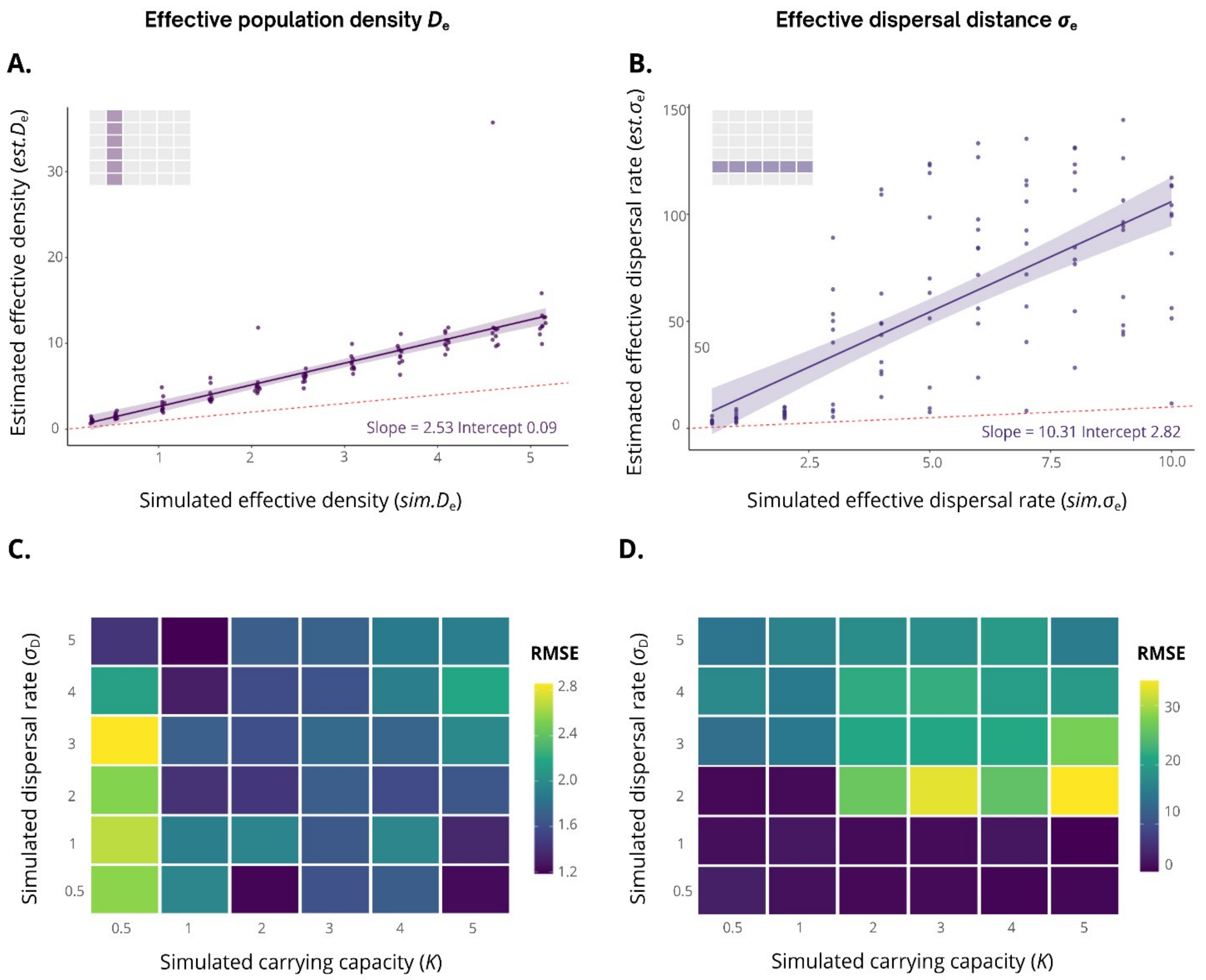


**Figure S3 - Accuracy of demographic parameter inference using *MAPS*.** Left panels: Effective population density ($D_e$). Right panels: Effective dispersal distance ($\sigma_e$). Effective density $D_e$ emerges from the simulated carrying capacity ($K$) and effective dispersal $\sigma_e$ emerges from the simulated dispersal distance ($\sigma_D$), linking simulation parameters to their emergent effective counterparts. **(A-B)** Relationship between simulated parameter values and their estimates inferred from IBD segments using *MAPS*. A dot represents one replicate; solid lines correspond to linear regressions between simulated and estimated values, with shaded areas indicating 95% confidence intervals. The dashed red line represents the 1:1 relationship. **(C-D)** Relative root mean squared error (RMSE) of parameter estimates across combinations of simulated carrying capacity ($K$) and dispersal distance ($\sigma_D$). Heatmaps illustrate how inference accuracy varies jointly with demographic conditions.

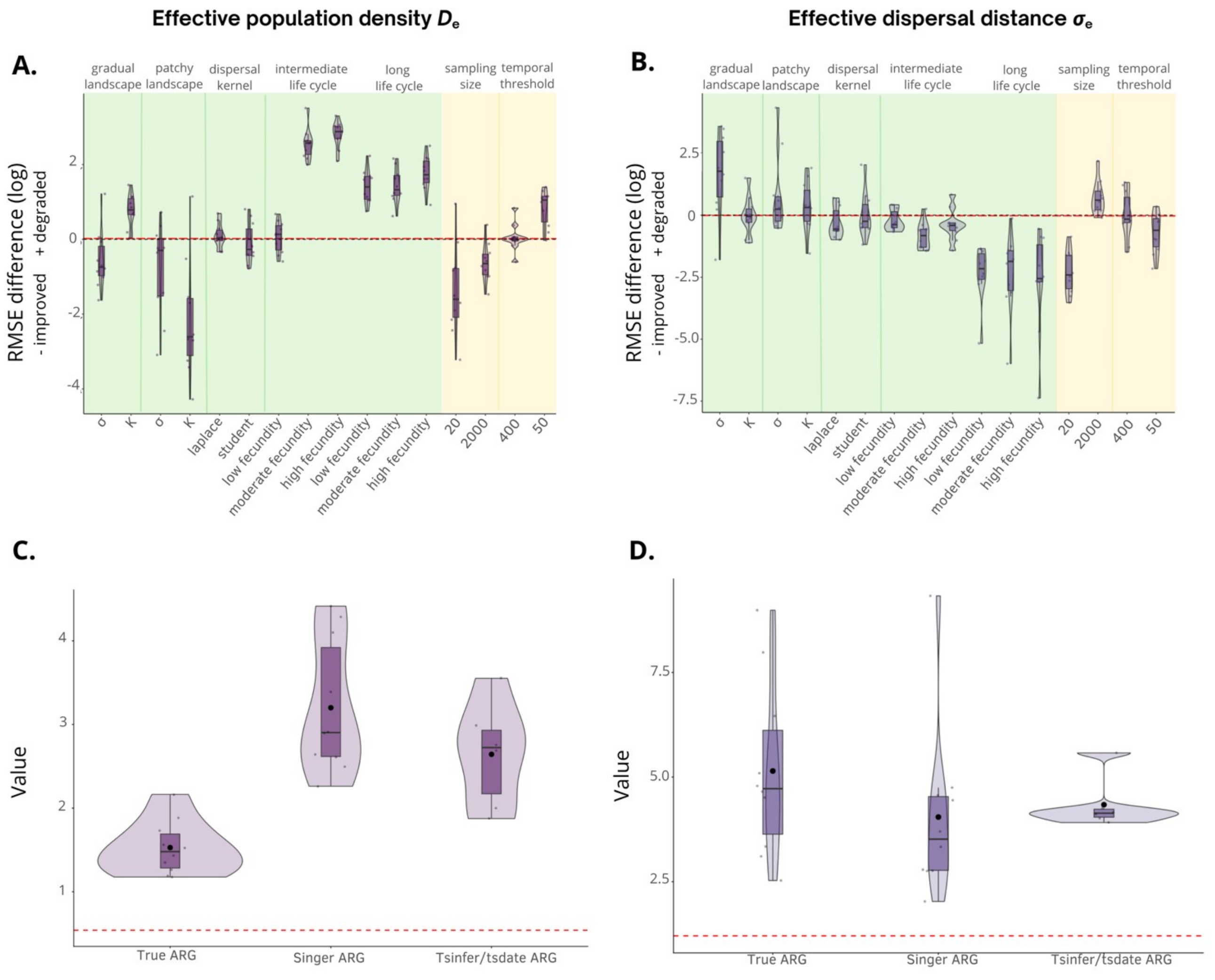


**Figure S4 - Robustness of parameter inference across tested conditions using MAPS.** Lefts panels: Effective population density ($D_e$). Right panels: Effective dispersal distance ($\sigma_e$). **(A-B)** Log-transformed relative root mean squared error (RMSE) of parameter estimates under each tested configuration compared with the reference configuration. Relative RMSE was calculated as the difference between the RMSE obtained under a given condition and that obtained under the reference configuration. The reference configuration is defined as average conditions: a homogeneous environment, a carrying capacity $K$=1, a dispersal variance $\sigma$=1, a gaussian dispersal kernel, a sample size of hundred individuals, a temporal depth of two hundred generations. The green area represents the ecological parameters, and the yellow area represents the methodological parameters. Positive value indicates a degradation of demographic inference relative to the reference condition. The red dashed line denotes no change relative to the reference configuration. **(C-D)** Value of parameters estimates obtained under different ARG reconstruction methods: true ARG, Singer ARG and tsinfer/tsdate ARG. The red dashed line represents the true parameter value.
